# *In-silico* Drug Repurposing Pipeline for Epilepsy: Integrating Deep Learning and Structure-based Approaches

**DOI:** 10.1101/2024.01.29.577686

**Authors:** Xiaoying Lv, Jia Wang, Ying Yuan, Lurong Pan, Jinjiang Guo

**Affiliations:** Global Health Drug Discovery Institute, Beijing, China; Cipher Gene Limited, Beijing, China; Ainnocence Limited, HongKong, China

**Keywords:** drug repurposing, epilepsy, gain-of-function gene, docking, deep learning, BBB (blood-brain barrier) permeability

## Abstract

Due to considerable global prevalence and high recurrence rate, the pursuit of effective new medication for epilepsy treatment remains an urgent and significant challenge. Drug repurposing emerges as a cost-effective and efficient strategy to combat this disorder. This study leverages the transformer-based deep learning methods coupled with molecular binding affinity calculation to develop a novel *in-silico* drug repurposing pipeline for epilepsy. The number of candidate inhibitors against 24 target proteins encoded by gain-of-function (GOF) genes implicated in epileptogenesis ranged from zero to several hundreds. Our pipeline has repurposed the medications with most anti-epileptic drugs (AEDs) and nearly half psychiatric medications, highlighting the effectiveness of our pipeline. Furthermore, Lomitapide, a cholesterol-lowering drug, first emerged as particularly noteworthy, exhibiting high binding affinity for 10 targets and verified by molecular dynamics (MD) simulation and mechanism analysis. These findings provided a novel perspective on therapeutic strategies for other central nervous system (CNS) disease.

## Introduction

Epilepsy, a central neurological disease, can be caused by any direct or indirect insult to the CNS, and often accompanied by cognitive and psychological impact due to its recurrent nature^1^. A meta-analysis study revealed that worldwide lifetime prevalence of epilepsy was approximately 1,099 per 100,000 individuals worldwide^2^. Recent treatment strategies for epilepsy primarily focused on the relief of symptoms, and advancements in treatment have enabled most patients to achieve seizure control^3^. Beside AEDs, alternative treatments such as dietary therapies^4^, surgery^5^, neuromodulation therapies^6^ can also aid in seizures control. Despite therapeutic advancements, reducing high recurrence rate for intractable epilepsy still poses a significant challenge for approximately 30% of patients^7^, even with monotherapy or polytherapy AED regimens^8,9^. Thus, the discovery of new AEDs would serve for not only improving seizure management but also for diminishing the substantial societal and healthcare burdens associated with these neurological conditions^10^.

Drug repurposing, the practice of finding new therapeutic usages for FDA-approved drugs, provides a cost-effective and time-efficient approach in drug discovery. Meanwhile, the integration of structure-based and deep learning methods has significantly enhanced drug repurposing, particularly in elucidating structure-activity relationships and accelerating drug discovery^11,12^. The Experimental Epilepsies (PDE3) database, developed by Nasir group through literature mining, has cataloged drugs that have demonstrated efficacy in experimental epilepsy models^13^. Reutens’s laboratory had attempted to repurpose drugs related to complement system for post-traumatic epilepsy, leveraging shared mechanism of action between these drugs and neuroinflammatory responses associated with epileptogenesis^14^. Moreover, potential epilepsy treatments have been identified by contrasting gene expression profiles in epileptic patients against healthy controls, using drug perturbation data from the Connectivity Map (CMAP) database^15^. A successful example of drug repurposing is Igalmi, initially approved by FDA in 1999 for sedation and analgesia in intensive care settings. BioXcel biopharmaceuticals repurposed it for epilepsy treatment, capitalizing on its properties as a selective α2-adrenergic receptor modulator known to alleviate agitation in adults^16^.

In this study, we developed an innovative *in-silico* drug repurposing pipeline by integrating structure-based and deep learning approaches. The pipeline is specifically designed to identify new potential inhibitors for epilepsy (Fig. 1). We selected 24 targets encoded by GOF genes for drug repurposing, restricted by the unavailability of crystal structures for other targets^17,18,19^. The identified potential inhibitors displayed a high BBB permeable probability to effectively reach brain tissue. We prioritized candidates that exhibited low cytotoxicity, especially those intended for chronic management. Consequently, our pipeline selected a range of drugs which may demonstrated anti-seizure activities against specific targets. Moreover, for Lomitapide repurposed by our pipeline, 100-ns MD simulations against each selected target and mechanism analysis were conducted to further validate its potential inhibition for epilepsy.

**Fig 1.**
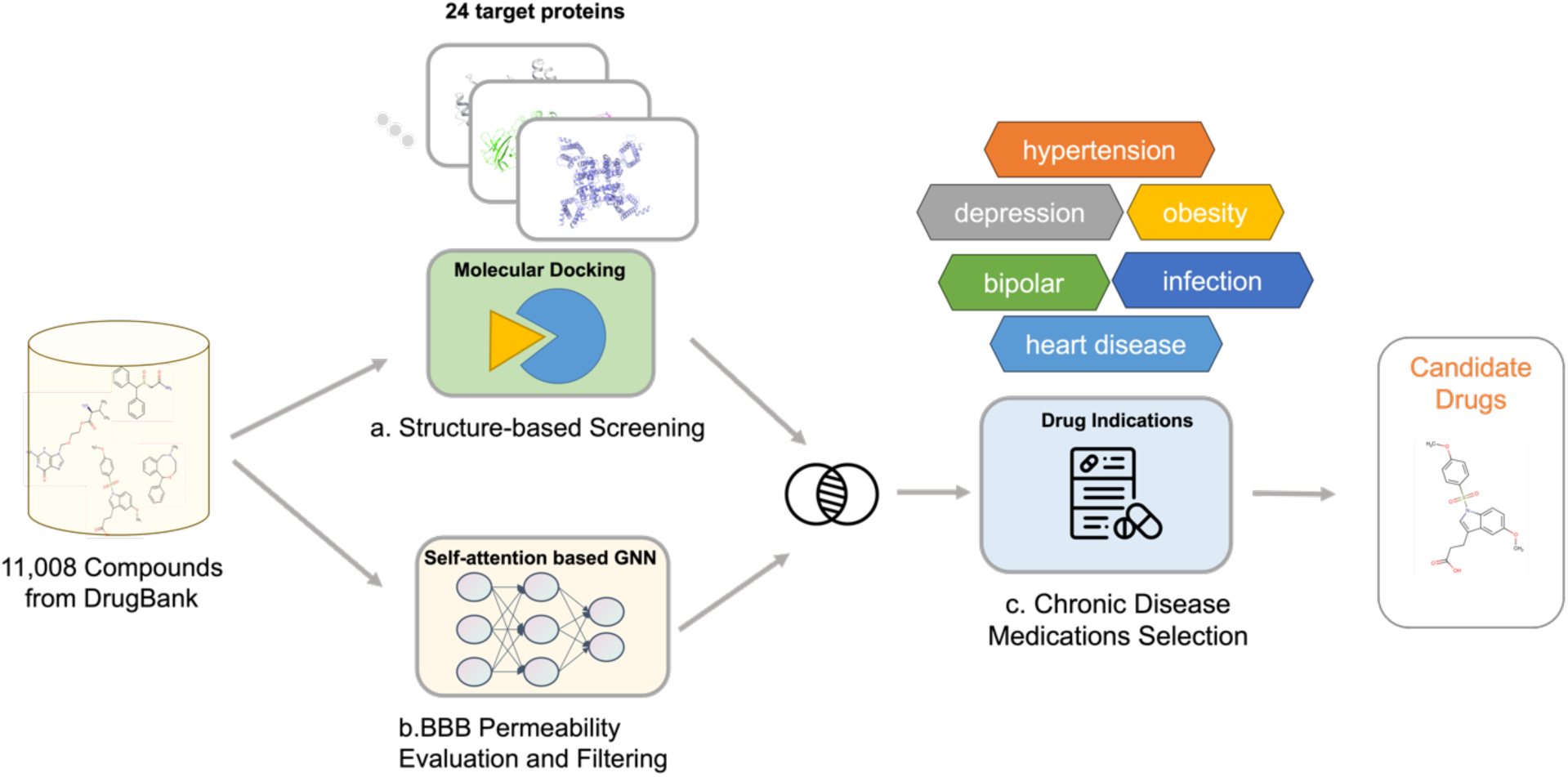
The *in-silico* drug repurposing pipeline for epilepsy. In our pipeline, we integrate two stages: structure-based screening (a) and BBB permeability evaluation and filtering (b) to screen potential medications from 11,008 compounds sourced from DrugBank database. (c) Medications prescribed for chronic diseases, including hypertensin, depression and infection disease, are preferred for seizure control in our *in-silico* pipeline.

## Results

### 2.1 Candidates number reduction by using individualized reference compounds

In the preliminary stage of our pipeline development, Glide standard precision (SP) docking was utilized to screen a set of 11,008 compounds against 24 target proteins encoded by GOF genes. When applying a uniformly established docking score threshold of -7.0, a highly disproportionate distribution of compounds meeting the criterion was observed across different targets (Fig. S1). To rectify this discrepancy, individualized docking score cutoffs were proposed for targets with a ligand-bound structure in the PDB database or known drug association. Notably, most reference compounds achieved docking scores below -7.0. Nevertheless, five reference compounds deviated from this trend, as detailed in Table 1. Remarkably, the *RHEB* encoding target, which has two distinct structures (PDB IDs: 6BT0, 6BSX), manifested docking scores for the non-GDP binding pocket of -6.08642 and -6.71108, respectively. When the individualized reference compounds were utilized as positive controls, a notable decrease in the pool of candidate compounds observed.

**Table 1.**
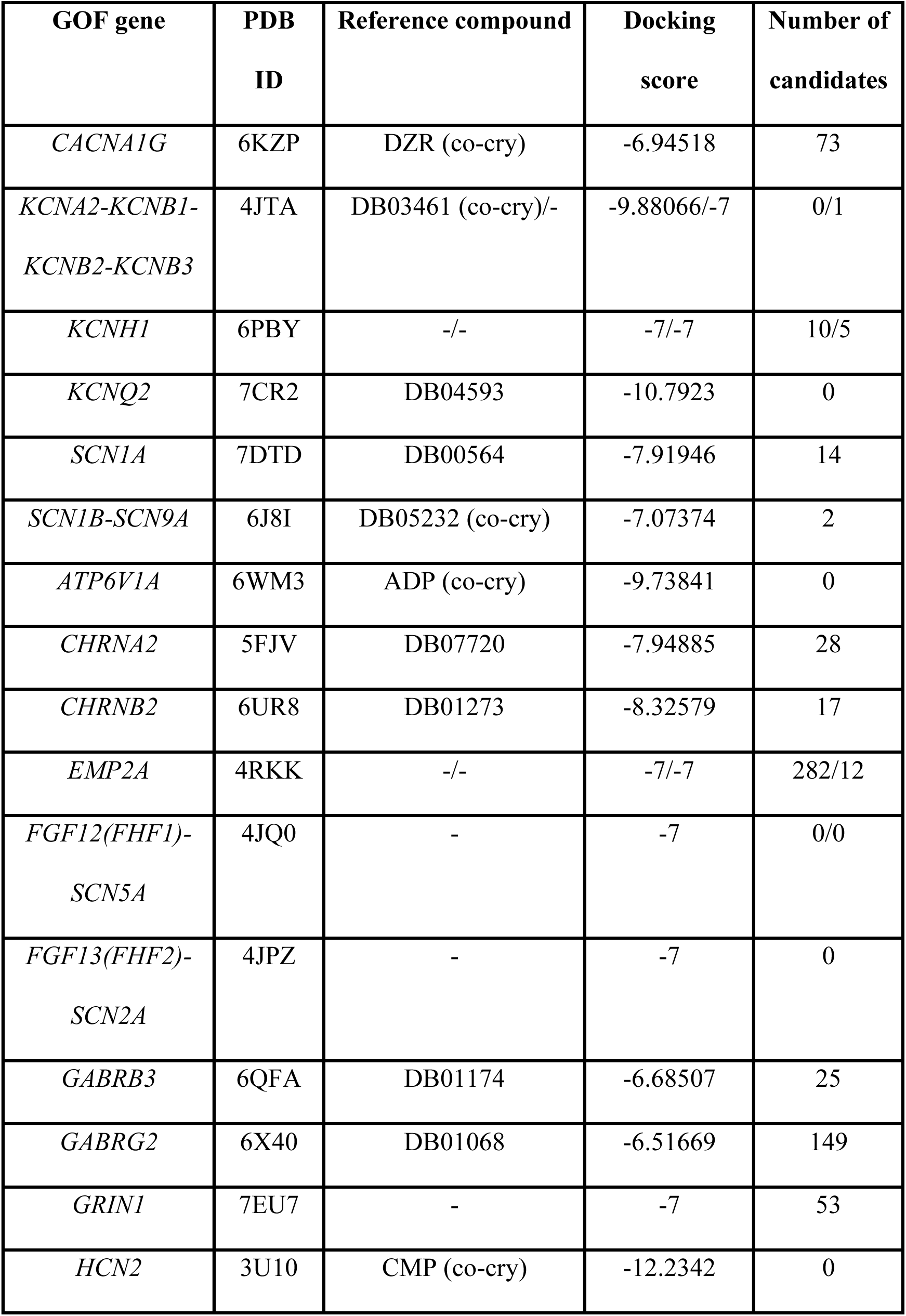
24 GOF targets and the number of repurposable drugs in our pipeline.

In the case of seven structures featuring two putative binding pockets, the quantity of compounds exhibiting strong binding affinity was generally higher in one pocket compared to the other. For example, when applying a cutoff of -7, the number of compounds with favorable binding affinity to the two pockets of protein (PDB ID: 4JTA) was 1637 for the first and 193 for the second (Fig. S1).

### 2.2 Compound BBB Permeability Evaluation

BBB permeability is paramount in determining the efficacy of drugs for neurological disorders by dictating the accessibility of compounds to CNS. To evaluate compound’s BBB permeability, we adopted a self-attention-based Graph Neural Network (GNN) model, namely Ligandfomer^20^, to predict its potential brain accessibility. For a given compound, the model generates a probability score within a range of 0 to 1, where a higher score indicates greater likelihood of brain accessibility. The Ligandformer model was rigorously trained and validated on the DB3B dataset^21^. The model’s performance was assessed by various metrics, yielding a harmonic average score of 0.932 for the area under the receiver operating characteristic curve (AUROC) and a harmonic average score of 0.925 for the area under the precision-recall curve (AUPR).

In our analysis of the predicted probability scores for 11,008 small molecule drugs, the distribution across various BBB permeability intervals appeared relatively uniform, with each interval containing approximately 1,000 compounds, except in the score ranges below 0.1 and above 0.9 (Fig. S2). Notably, the BBB permeability predictions for AEDs, which are commonly available in the market, exceeded a probability of 0.8 (Table S1). The consistent pattern underscores the model’s robust capability for predicting BBB permeability. To avoid omitting any potential drugs for epilepsy, we selected 3,200 candidate drugs with BBB permeable probability score above 0.75 for the subsequent stage.

### 2.3 The potential drugs screened against each target

In our strategic pipeline, there was a preferential selection for the orally administered and FDA-approved drugs for chronic diseases. We identified a multitude of prospective compounds for most targets. However, for eight target proteins encoded by—namely, *KCNQ2*, *ATP6V1A*, *FGF12*, *FGF13*, *HCN2*, *RHEB* and *SCN8A* genes—there were no suitable potential medications available, as detailed in Table 1. Despite the compatibility of numerous drugs with their target crystal structures in the docking analysis, none exhibited sufficient probabilities of BBB permeability, based on the Ligandformer predictions.

In the joint filtering analysis, the number of potential candidate medications range from zero to several hundreds, particularly exceeding one hundred for the target (pocket 1) encoded by *EMP2A* and *GAGRB2* genes (Fig 2). The lack of a reference compound for target encoded by *EMP2A* likely contributed to the substantial number of potential drugs. Nearly half of candidate drugs are indicated for psychiatric conditions, such as depression, schizophrenia, insomnia, bipolar, Parkinson’s disease, dementia, anxiety and migraine, all of which display symptoms and mechanisms closely associated with the onset of seizures. Consequently, these drugs are deemed promising for the prophylaxis of seizure incidents. Furthermore, our pipeline identified 27 antiepileptic drugs (AEDs), confirming its effectiveness in identifying suitable treatments for epilepsy. Additionally, a few antihypertensive and antidiabetic drugs were recognized for their potential efficacy in epilepsy management (Table S2).

**Fig 2.**
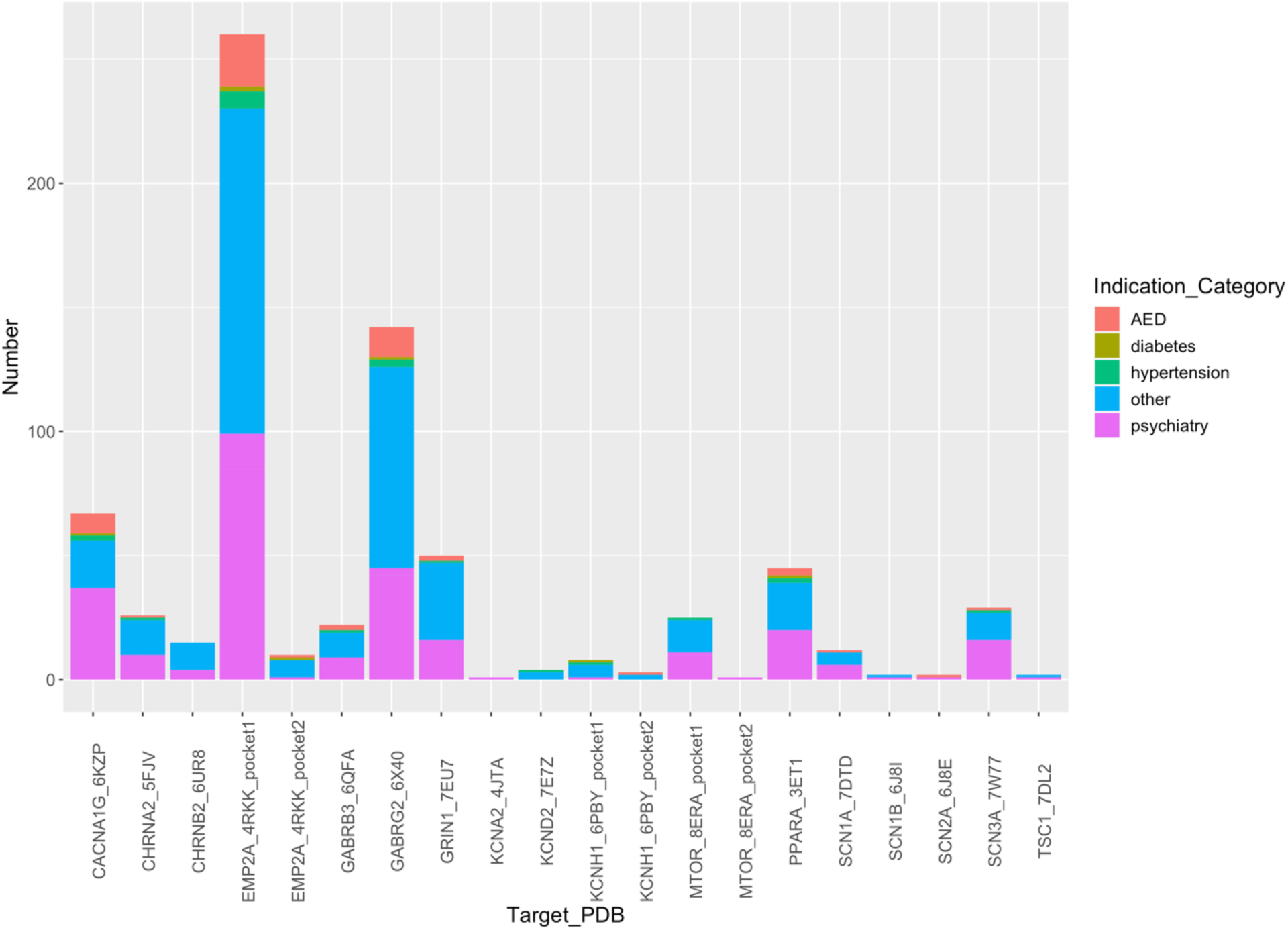
The distribution of candidate drugs selected by our *in-silico* pipeline. AED means anti-epileptic drug.

In our investigation of voltage-gated sodium channels, we analyzed five structures (encoded by *SCN1-3A, SCN8-9A*). Despite the structures’ high degree of similarity, indicated by a root mean square deviation (RMSD) of less than 2 Å (Fig. S3a), distinct binding sites were identified (Fig. S3b). This dissimilarity in binding sites could explain the observed inconsistency in the efficacy of candidate drugs targeting the SCN family (Table S2). For the *SCN1A* encoding target, Midazolam was recognized for its ability to generate π-π interactions with Phe1789 (Fig. 3a), exhibiting a notable docking score of -8.21737. Fluphenazine (Fig. 3b) and Lomitapide (Fig. 3c) emerged as the most promising drugs for target encoded by *SCN1A* gene, offering multiple protein-ligand interactions that enhance pose stability.

**Fig 3.**
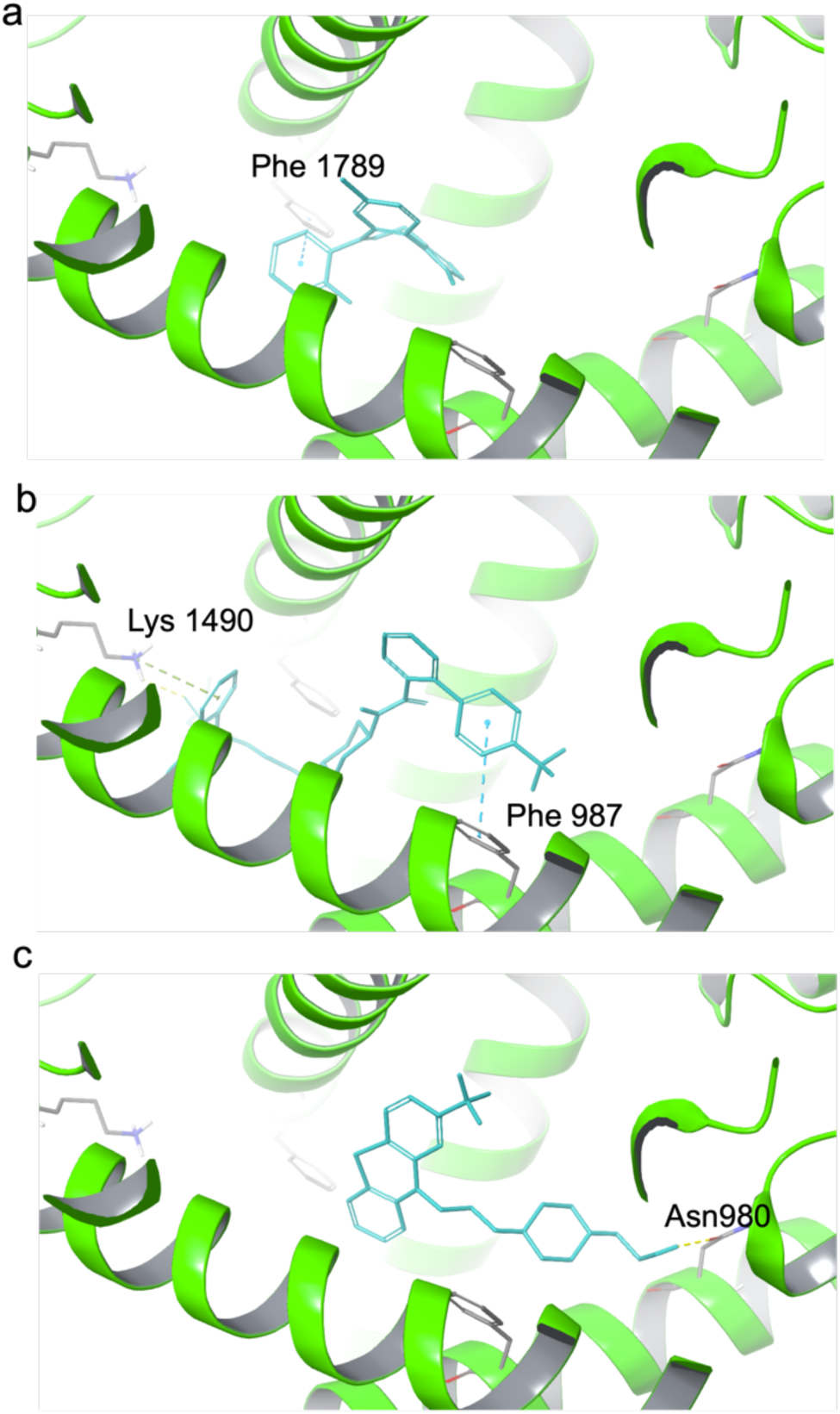
The binding pose of Midazolam, Lomitapide and Fluphenazine in SCN1A target. The 7DTD protein (cartoon) is colored by green and the ligands (stick) (a: Midazolam, b: Lomitapide, c: Fluphenazine) by cyan. The key residues (stick) in protein are colored by gray. The hydrogen-bond, cation-pi interaction and pi-pi interaction are colored by yellow, green and blue dotted line, respectively.

### 2.4 Potential medications targeting multiple proteins

In this study, several drugs were identified as candidates for repurposing with actions on multiple targets implicated in seizure control. Among the identified drugs, Nebivolol, Lomitapide, and Pimozide exhibited the most promise, engaging with 11, 10 and 9 targets respectively (Table 2).

**Table 2.**
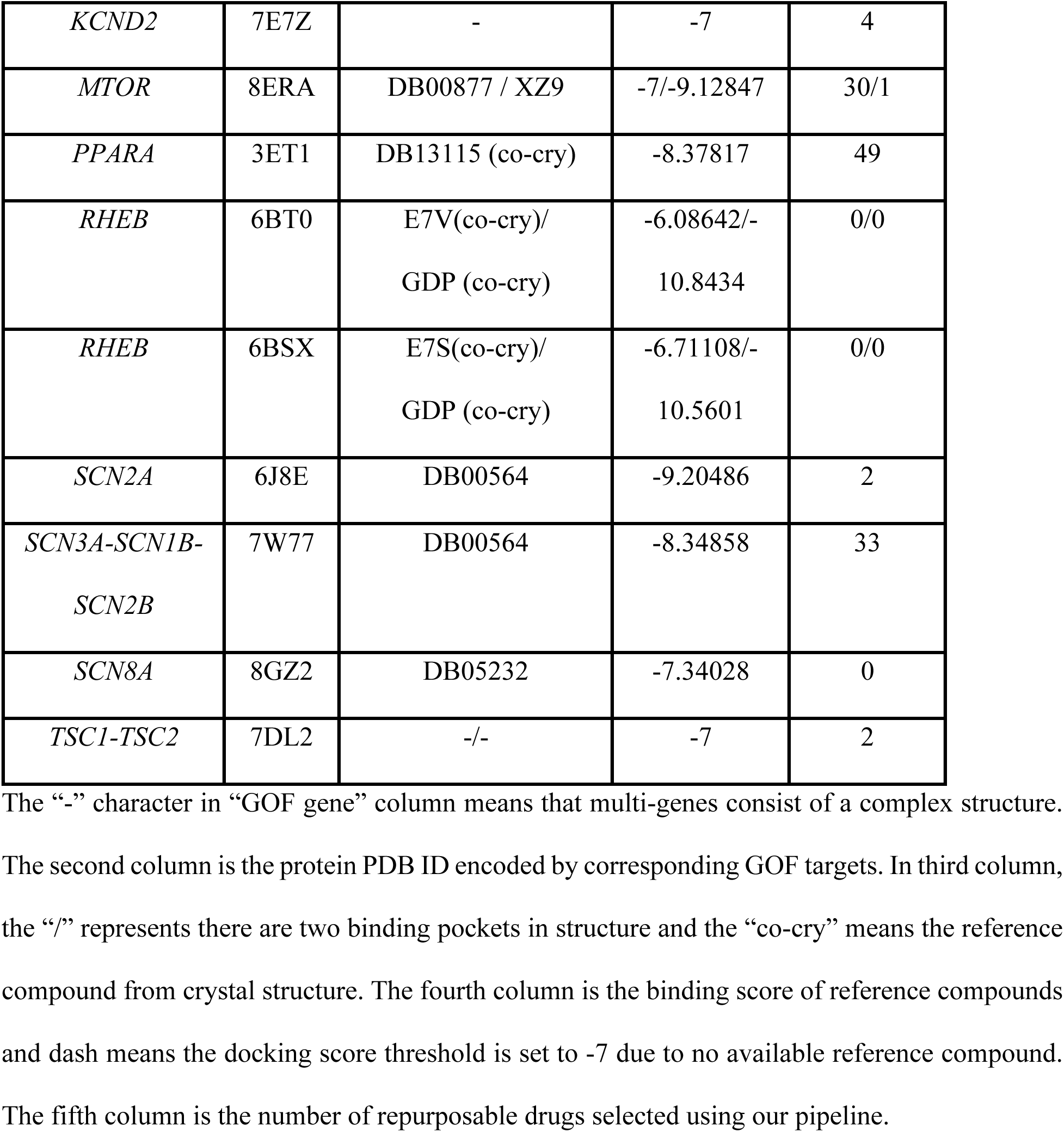
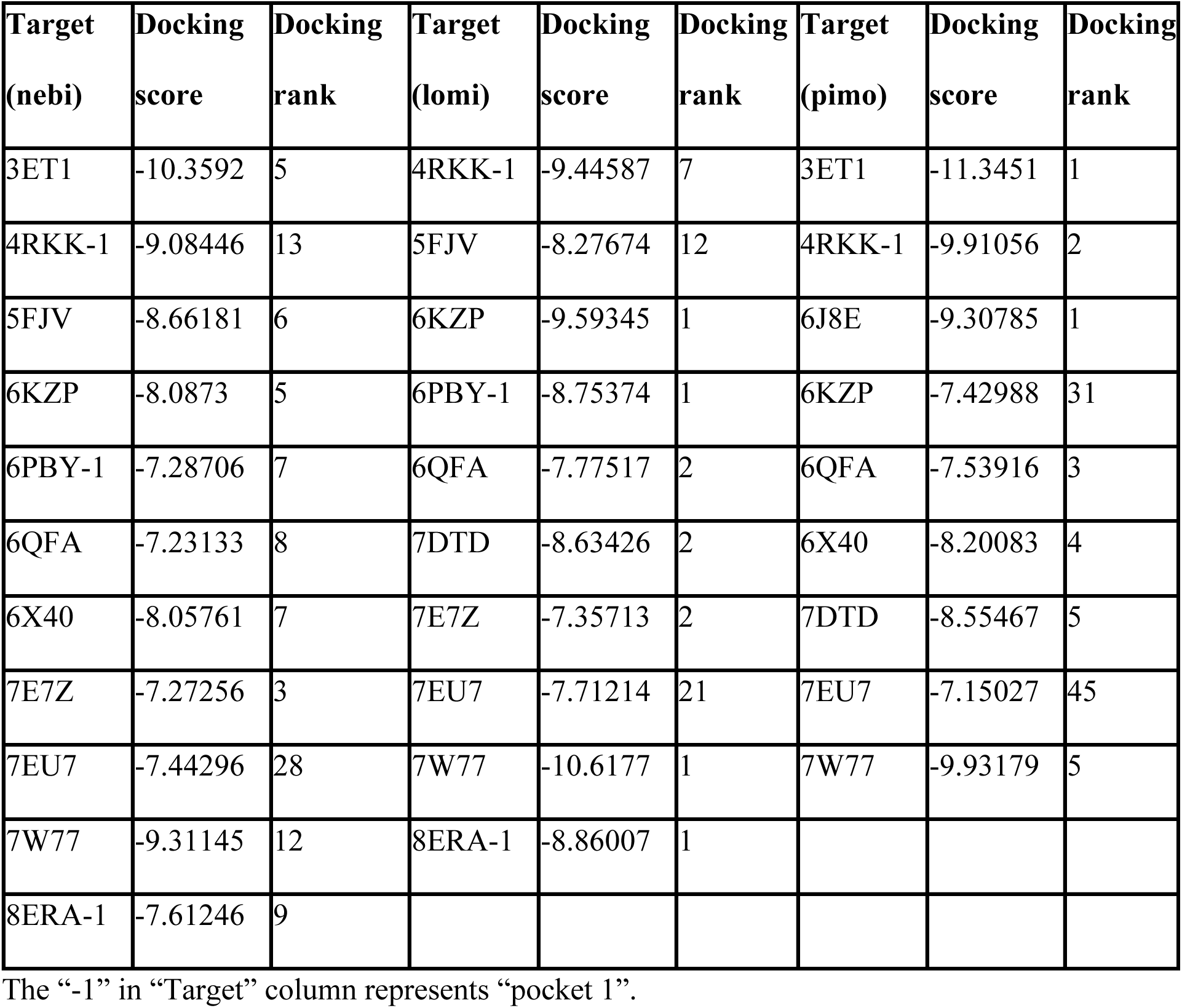
The docking score and rank of Nebivolol, Lomitapide and Pimozide in drug list.

Nebivolol was a selective β1-blocker with vasodilatory properties commonly prescribed. Extensive research endeavors have sought to unravel the intricacies of drug-drug interactions between nebivolol and prevalent adverse drug events using a murine model^22^. Additionally, a synergistic effect is observed when nebivolol is combined with a low dose of phenytoin, effectively mitigating seizures precipitated by maximal electroshock seizures (MES)^23^. Conversely, nebivolol has been demonstrated to diminish the anticonvulsant efficacy of both carbamazepine and phenobarbital in MES models^24^.

Lomitapide, utilized for cholesterol reduction in patients with homozygous familial hypercholesterolemia, has emerged as a candidate with potential efficacy for 10 distinct targets (Fig. 4a). Despite its demonstrated affinity for multiple GOF targets, Lomitapide lacks documented evidence in treating neurological disorders. The stability of the protein-ligand interaction in *CACNA1G* encoding target (PDB ID: 6KZP), is largely due to a π-π interaction with Phe917 and a hydrophobic interaction, as depicted in Figure 4b. Comparable interactions are seen with the target encoded by *MTOR* gene, where Lomitapide binds similarly through π-π and hydrophobic interactions (Fig. 4c). Moreover, the significant docking score of -10.6177 for the target encoded by *SCN3A* gene is primarily the result of hydrophobic interactions, highlighted in Figure 4d.

**Fig 4.**
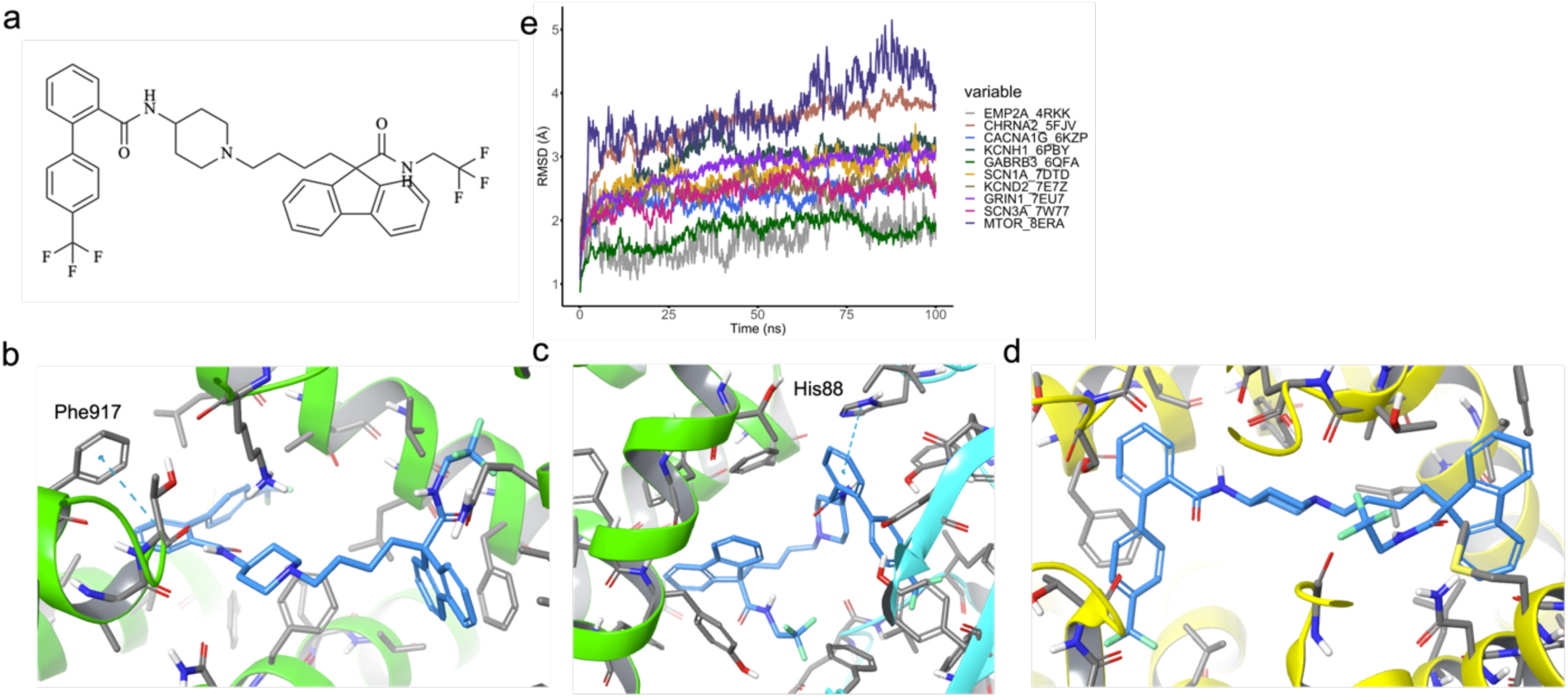
Lomitapide structure and binding pose in selected multiple targets with protein backbone RMSD in 100-ns MD simulations. The 2D structure of Lomitapide is in panel a. The binding poses of Lomitapide in CACNA1G (panel b), MTOR (panel c) and SCN3A (panel d) targets. These targets are colored by chain name (chain A: green, chain B: blue, chain D: yellow). The key residues and ligands (stick) in protein are colored by gray and blue, respectively. The pi-pi interaction is colored by blue dotted line. The RMSD of protein backbone (referred by initial docking pose) for 100-ns MD simulation in each target are showed in panel e.

Pimozide, associated with nine targets as indicated in Table 2, is a potent dopamine antagonist. Studies have shown this medication to be correlated with a reduced risk of inducing seizures^25,26^.

### 2.5 MD simulation for Lomitapide stability validation against 10 selected targets

To further substantiate the potential of Lomitapide in mitigating epilepsy progression, MD simulations were conducted for each target. Stability of the ligand-binding domains was observed in all 10 systems, as evidenced by convergence of backbone RMSD within the 100 ns production phase (Fig. 4e), with the exception of the proteins encoded by *CHRNA2* and *MTOR* genes. For the *CHRNA2* encoding target (PDB ID: 5FJV), a PDB resolution of 3.2 Å suggests that suboptimal side-chain packing may account for the observed instability during MD simulation. In the case of target encoded by *MTOR* gene (PDB ID: 8ERA), the elevated RMSD was primarily attributed to the relative movement of chains A and B, yet the RMSDs of these individual chains remained approximately 2 Å (Fig. S4a).

In most systems, Lomitapide maintained a pose similar to that of the initial docking conformation, with only minor deviations observed within the binding pocket during the 100 ns MD simulations, as depicted in Movies S1-S10. Stability of the ligand within the binding site was observed, with the notable exception of the ligand segments located in the solvent or cavity, such as those in 4RKK and 5FJV, which exhibited flexibility due to the lack of direct interactions with the receptor. In the sirolimus binding pocket, situated at the protein-protein interface of the *TSC* complex (PDB ID: 8ERA), the two chains engaged by sirolimus binding showed significant separation, potentially resulting from the weaker interactions of Lomitapide compared to the macrocyclic lactones sirolimus. Despite Lomitapide’s high docking score, the 5 hydrogen bonds with Asp38, Gln54, Glu55, Ile57 and Tyr83 of protein domain (chain B of 8ERA) encoded by *FKBP1A* gene were not present (Fig. S4b). Additionally, the new pi-pi interaction with His88 in *FKBP1A* encoding protein proved insufficient to sustain the interaction between chain A and chain B (Fig. S4c).

### 2.6 Mechanism of action exploration of Lomitapide

To further explore Lomitapide’s mechanism of action, we investigated its target’s interacting genes within a protein-protein interaction (PPI) database and searched for shared pathways within AEDs in Gene Ontology (GO) database. The reported target of Lomitapide in FDA is microsomal triglyceride transfer protein, encoded by *MTTP* gene. In our curated PPI database^27^, six genes (*APOA1*, *APOA2*, *APOA4*, *APOB*, *P4HB*, *HSP90B1*) demonstrated interactions with the *MTTP* gene. The majority are apolipoprotein-related gene, with the exception of the *HSP90B1* gene (Fig 5a), which has been previously associated with diseases such as Pelizaeus-Merzbacher-Like disease and bipolor disorder^28^. Notably, the former, Pelizaeus-Merzbacher-Like disease type 1, is a hereditary central nervous system disorder that exhibits several symptoms reminiscent of epilepsy.

**Fig 5.**
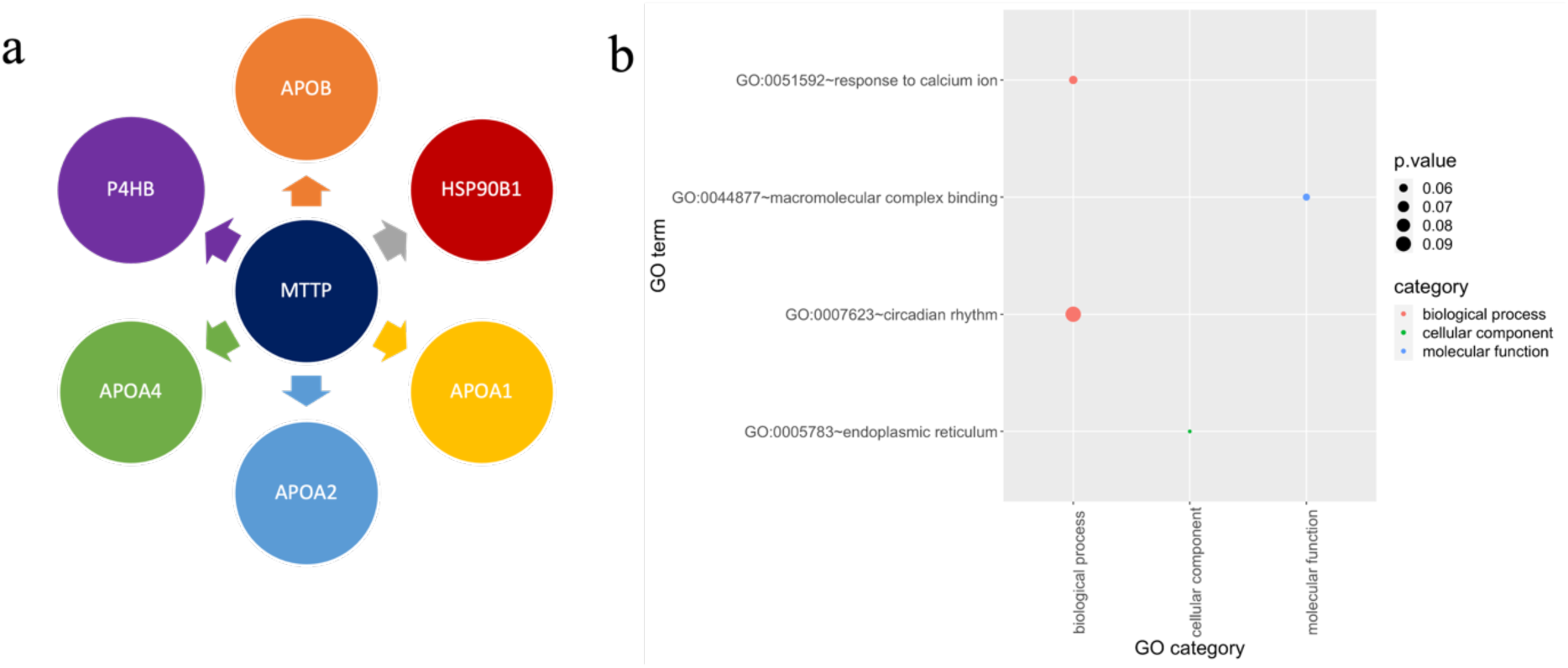
The mechanism of action exploration of Lomitapide. The genes that interact with MTTP gene in PPI database are shown in panel a. Panel b is the shared pathway of targets between Lomitapide and AEDs. The size of cycle represents p-values and the GO categories being distinguished by color: red for biological process, green for cellular component and blue for molecular function.

We conducted a search for pathways related to Lomitapide in the GO database, identifying 41 pertinent pathways (Table S3). Subsequently, we enriched the target genes of AEDs to analyze pathways using the DAVID website^29^. Our analysis identified four shared pathways: response to calcium ions, macromolecular complex binding, circadian rhythm regulation, and endoplasmic reticulum (Fig 5b). These findings suggest the potential existence of a common mechanism of action for Lomitapide and AEDs.

## Discussion

Several prior studies on drug repurposing for epilepsy have primarily utilized approaches such as literature search^13^, drug-target network^7^ and gene expression profiling^15^, as previous mentioned. In contrast, our proposed pipeline aims to repurpose drugs for seizure management by integrating structure-based and deep learning methods. Since FDA-approved drugs already conform to established clinical standards of safety and bioavailability, our focus is primarily on evaluating their on-target binding affinity and CNS bioavailability.

Our *in-silico* pipeline has effectively identified numerous potential therapeutics; nonetheless, it is subject to inherent constraints of the approaches employed in our study. The accuracy of docking scores depends on various factors, including, but not limited to, the resolution of the receptor, the conformation of the ligand, and the selected scoring function. On the other hand, the deep learning model for predicting BBB permeability demonstrated promising performance, however its predictive accuracy is still limited by the quality and diversity of training data and inherent algorithmic constraints. Therefore, it is essential to validate these computational predictions with experimental edassays.

In conclusion, we have established an *in-silico* drug repurposing pipeline, leveraging structural compatibility and deep learning method to screen FDA-approved drugs for potential inhibitory effects on targets encoded by 24 GOF genes implicated in epilepsy. We observed a considerable variation in the number of potential repurposable drugs, which ranged from zero to several hundreds, contingent on the reference compounds used for each target. Notably, our analysis identified common AEDs as possible candidates, underscoring the effectiveness and efficiency of our pipeline in drug repurposing efforts. For the first time, Lomitapide emerged as a promising candidate medication, demonstrating stability across 10 targets in MD simulations and shared similar mechanism with AEDs in PPI interaction and GO enrichment analysis. This finding warrants further investigation into its therapeutic potential using zebrafish models and subsequent validation in human patients. While our approach offers insightful directions for epilepsy therapy, extensive research remains imperative to realize the ultimate objective of curing this complex disease.

## Materials and Methods

### 4.1 GOF genes searched from three databases

Potential candidate genes implicated in epileptogenesis via GOF mechanisms were identified through database search in PubMed Central^30^, Online Mendelian Inheritance in Man (OMIM)^31^ and Development Disorder Genotype - Phenotype Database (DDG2P) using “epilepsy” and “gain of function” as keywords. Finally, 72 genes were selected as target genes for drug repurposing. The reported evidence for GOF spans three levels-variant, gene, disease-detailed in Supplemental Table S4.

### 4.2 Crystal structures on target proteins and small molecules

Given that certain proteins function as complexes, a single crystal structure can suffice for multiple genes. Only 25 available crystal structures encoded by GOF genes with appropriate binding pockets were downloaded from the PDB database^32^ for the repurposing of small molecules to enhance docking precision (Table 1). Specially for the *RHEB* encoding target, two distinct crystal structures were selected due to a significant conformational alteration at the Tyr74 amino acid, as illustrated in structures 6BT0 and 6BSX (see Fig. S5). From the DrugBank database (version *2022.1.17*)^33^, we imported 11,008 small molecules for the purpose of docking with the targets.

### 4.3 Structure-based molecule docking

The docking workflow encompassed three steps: protein preparation, ligand preparation and the docking operation. Protein preparation utilized schrodinger Protein Preparation Workflow panel^34^ with default parameters via Maestro interface (version *12.9.123*, released at 2021.3). This process entailed the designation of bond orders, addition of hydrogen atoms, retention of waters beyond 5Å distance with ligands, and the construction of missing side chains and loops. Subsequently, H-bond assignments were optimized based on PROPKA-predicted pKa values, followed by a restrained energy minimization using the default force field.

The putative binding pocket was defined either by referring the position of co-crystallized ligands or through prediction utilizing the SiteMap suite^35^. While most proteins possessed a single binding pocket, six proteins presented two putative binding pockets (Table 1). Protein grid files were generated by Receptor Grid Generation panel. After ligand preparation using command: ligprep-ismi *.smi-omae *maegz, a total of 11,008 small molecules were successfully converted from 2D SMILES representations to 3D structures. These compounds were then docked into respective putative binding pockets employing the Glide SP docking protocol, which utilized the OPLS_2005 forcefield and default parameters^36,37^.

### 4.4 BBB Evaluation and Filtering

We utilized Ligandformer, a deep learning model based on transformer architecture to estimate BBB permeability. Ligandformer, as a graph neural network that employs a multi-layer, single-head self-attention mechanism to predict ligand properties. This model leverages the self-attention mechanism inherent in deep learning architecture, allowing a detailed representation of the relationship between chemical structures and their attributes. This capability facilitates more informed interpretations, as demonstrated in Figure S6.

The predictive model was trained and validated with a dataset comprising experimentally determined BBB properties, as reported by Meng et al^21^. This dataset contained 7,806 data points, which included 4,955 positive samples (permeable) and 2,851 negative samples (non-permeable). To enhance the model’s generalization capabilities, we employed a 5-fold cross-validation method. During each iteration of this process, the dataset was divided into an 80% training set and a 20% testing set, guaranteeing exposure to diverse data distributions. Candidate drugs approved by FDA for chronic conditions Any medications related to cancer treatment were excluded using keywords such as “malignant tumor”, “cancer”, “carcinoma” in their indication and their approved status was “true” in the DrugBank database.

### 4.5 System setup and MD simulations for 10 potential Lomitapide targets

We conducted MD simulations on the docking conformations produced by Lomitapide for 10 distinct targets. The initial structures were derived from the optimal docking conformations and maintained the same protonation state for the titratable residues. Given the substantial size of *GRIN1* encoding protein (PDB ID: 7EU7), MD simulations were confined to the transmembrane region alone (Fig. S7). We utilized the Tleap module in AMBER 22^38^ to systematically delete and subsequently reintroduce hydrogen atoms. For the three solvent proteins (PDB IDs: 4RKK, 5FJV, 8ERA) systems, we built a TIP3P water box along XYZ axis, extending 12 Å and neutralized the entire systems with 0.15 M NaCl concentration. For the rest seven membrane protein systems, we additionally oriented the direction of transmembrane region along Z-axis using PPM 2.0^39^ and embedded the proteins into 1-palmitoyl-2-oleoyl-glycero-3-phosphocholine (POPC) bilayer using CHARMM-GUI membrane builder^40^.

All-atom MD simulations were conducted employing PMEMD engine in AMBER 22^38^. The AMBER FF19SB force field^41^ and AMBER lipid force field LIPID21^42^ was adopted for protein and POPC molecules, respectively. Lomitapide was parameterized using general AMBER force field (GAFF). A cutoff of 10 Å was implemented for nonbonded interactions and bond-length constrain involving hydrogen atoms were managed by the SHAKE algorithm^43^. The particle mesh Ewald (PME) algorithm^44^ was used to address long-range electrostatic interactions. Simulations were conducted under periodic boundary conditions in all three dimensions. Initially, energy minimization was performed for 10,000 steps, followed by a gradual heating of the system from 0 K to 310 K across a span of 500 ps with the aid of the Langevin thermostat^45^. During heating, positional restraints were applied to the heavy atoms of the proteins and the ligand with a force constant of 50 kcal/mol/Å^2^. This was succeeded by 2 ns of pre-equilibration in the isochoric-isothermal (NVT) ensemble with the imposition of decreasing positional restraint force from 50, 5 to 0.5 kcal/mol/Å^2^. For seven membrane protein systems, the lipid heads were also subjected to positional restrained and a further 30 ns equilibration phase was administered with the proteins and ligand still under 0.5 kcal/mol/Å^2^ constraint. The production phase spanned 100 ns for each system in a constant pressure (NPT) ensemble. The time step was set to 2 fs. The frames were saved every 5,000 steps for trajectory analysis.

### 4.6 PPI interaction and Gene Enrichment analysis

The target gene *MTTP*, associated with Lomitapide, was searched for interacting genes in the PPI database established in our previous study^27^. We investigated pathways related to Lomitapide within the GO database^46,47^ (Table S3). Furthermore, target genes of AEDs were retrieved from the DrugBank database and analyzed for GO pathway enrichment using the DAVID platform ^48,29^, examining three GO categories (biological process, cellular component and molecular function) using default settings (Table S3).

## Author contributions

J.J.G carried out study design and performed the BBB permeability prediction. J.W. conducted GOF genes search and help to write the partial Introduction part. X.Y.L. handled the remaining part and write the manuscript. Y.Y. and L.R.P provided advice on study design and help to revise the manuscript. All authors reviewed the results and approved the final version of the manuscript.

---

**Fig S1.**
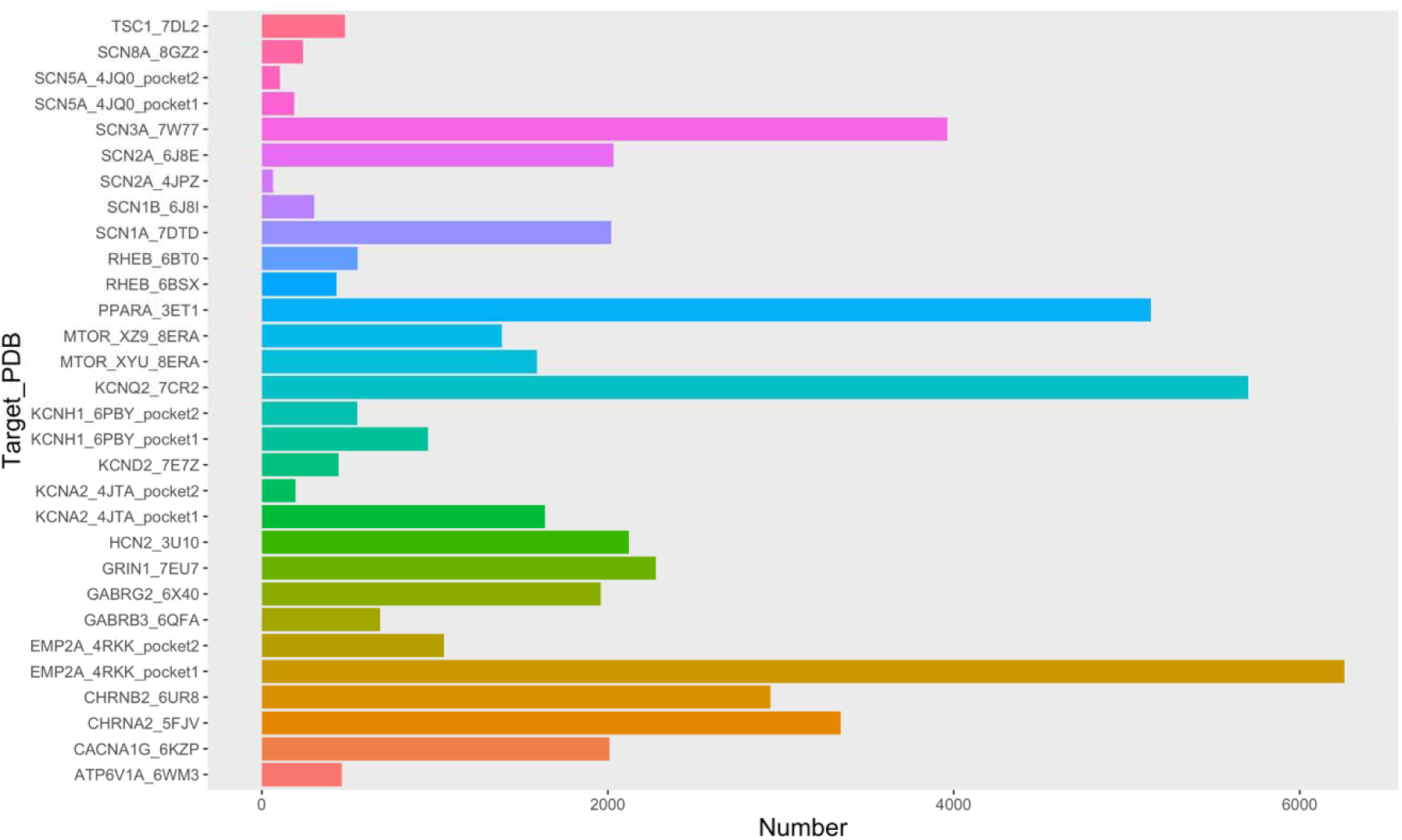
The number of candidates after docking filter using -7.0 as threshold criterion.

**Fig S2.**
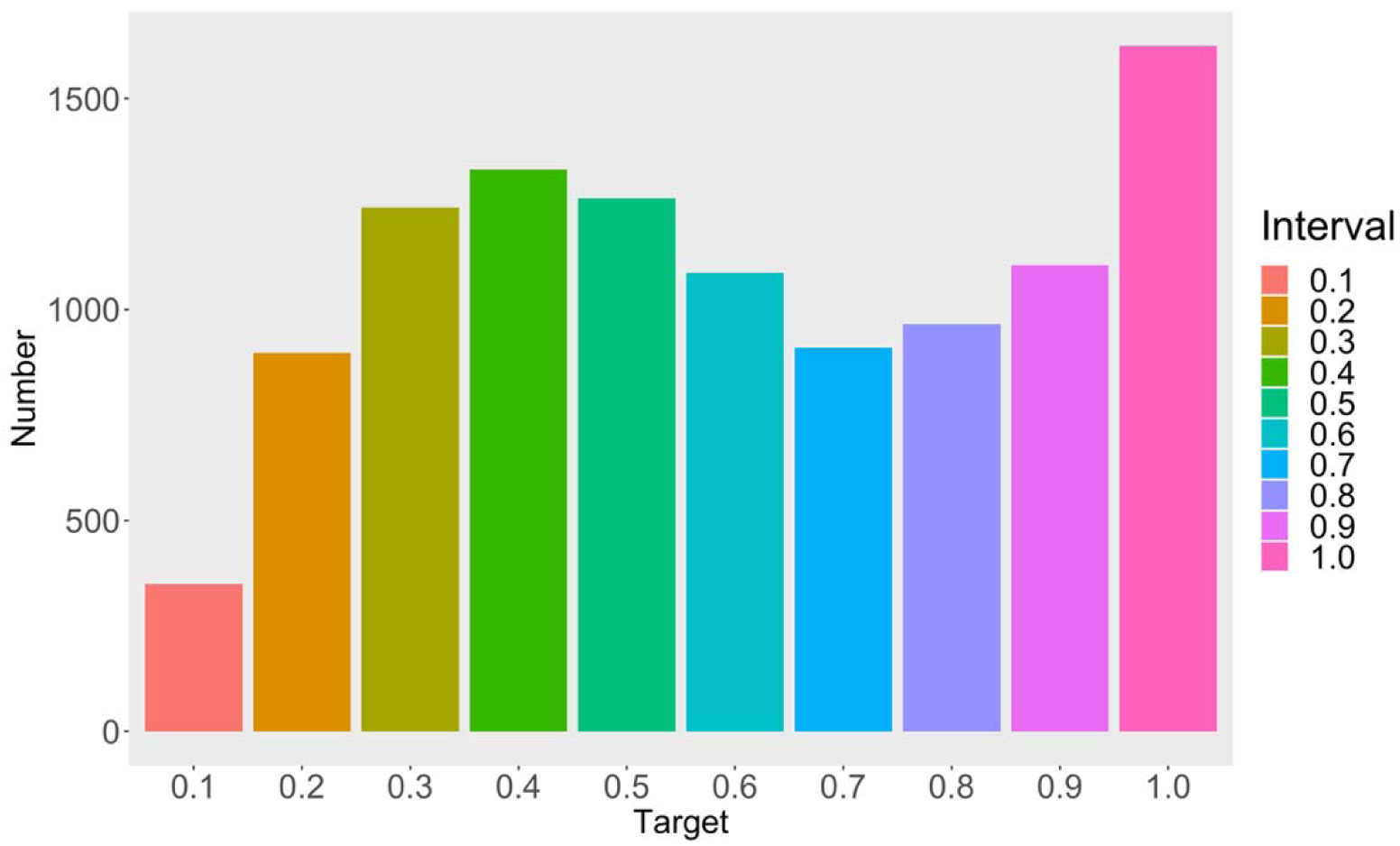
The distribution of predicted BBB permeability probability for compounds from DrugBank database.

**Fig S3.**
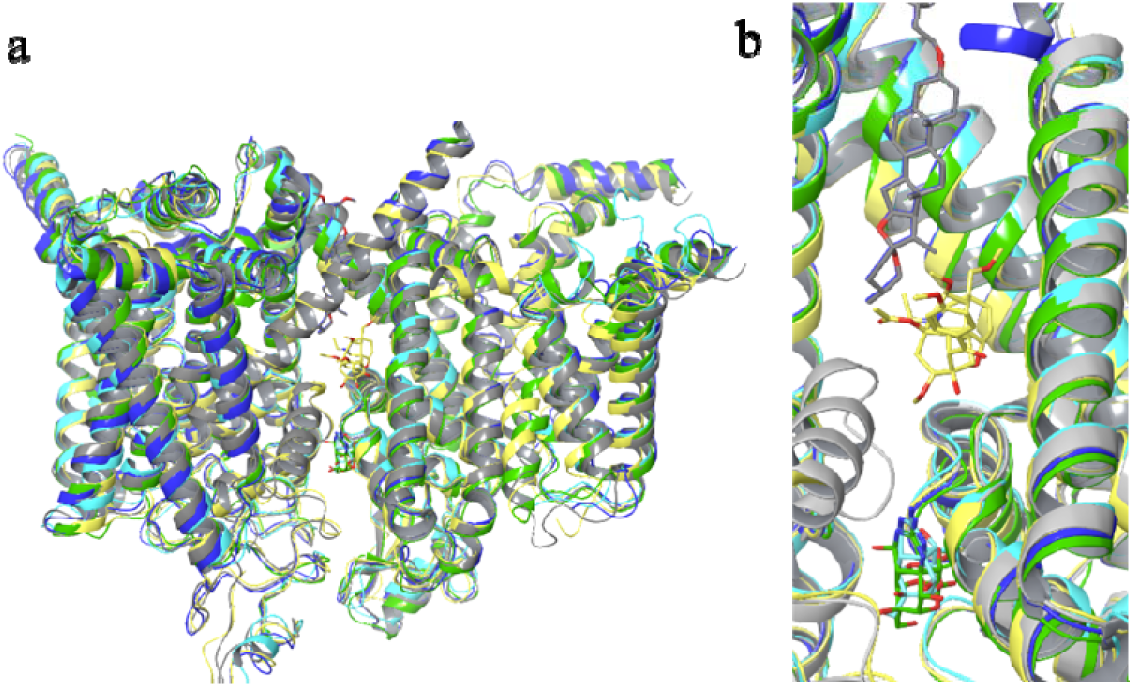
The five voltage-gated sodium channel structures. a. Using SCN1A (7DTD:gray) as a reference structure, the other four structures were aligned using protein (cartoon). The RMSD for SCN2A (6J8E:blue), SCN3A (7W77:yellow), SCN8A (8GZ2:cyan) and SCN9A (6J8I:green) are 1.664, 1.373, 1.779 and 1.664, respectively. b. The close-up view of panel a. These ligands are represented by stick with the same color of corresponding protein in panel a.

**Fig. S4.**
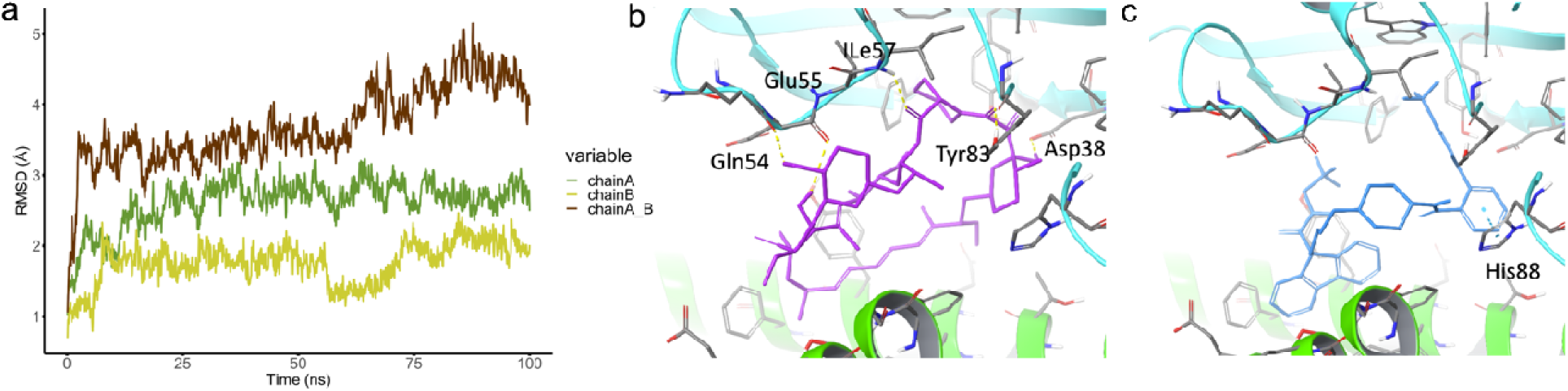
The RMSD plot and ligand-receptor interaction for the target (PDB ID: 8ERA). Panel a, RMSD plot for the protein backbone, calculated across 8ERA complex and its chain A and B during the 100ns production phase. In panel b and c, chain A encoded by *MTOR* gene and chain B encoded by *FKBP1A* gene was highlighted in green and cyan, respectively. The co-crystal XYU (panel b) and Lomitapide (panel c) were colored by purple and blue, respectively.

**Fig S5.**
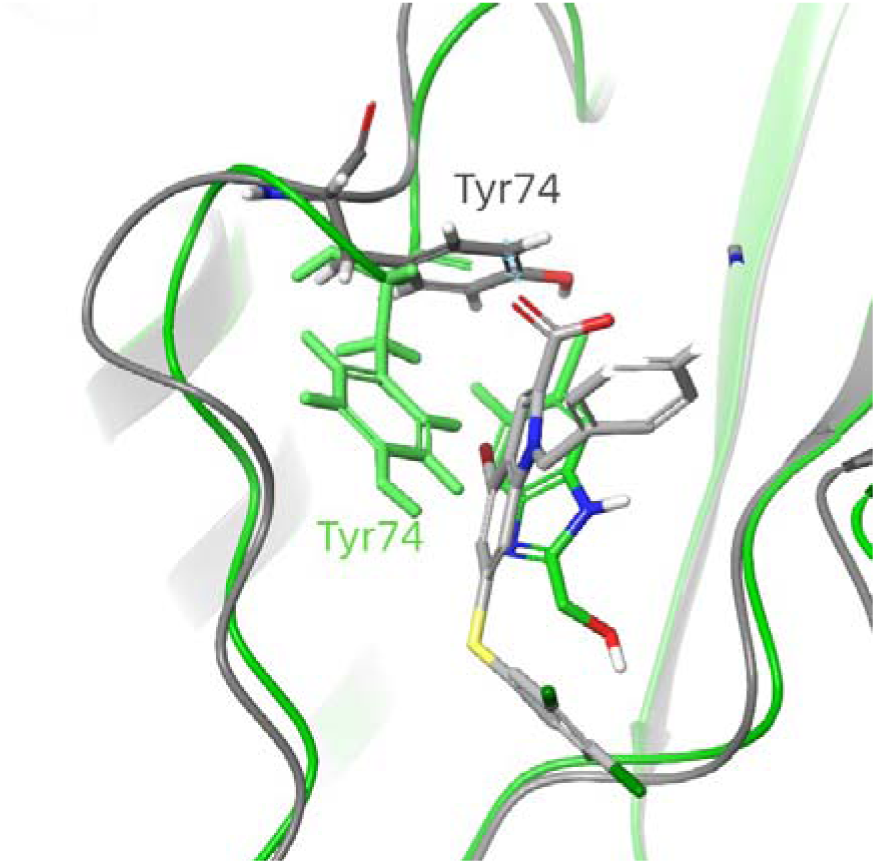
The two conformations of Tyr74 in 6BT0 (gray) and 6BSX (green) crystal structures. For RHEB target, the binding poses of ligand E7V (gray) and E7S (green) are shown.

**Fig S6.**
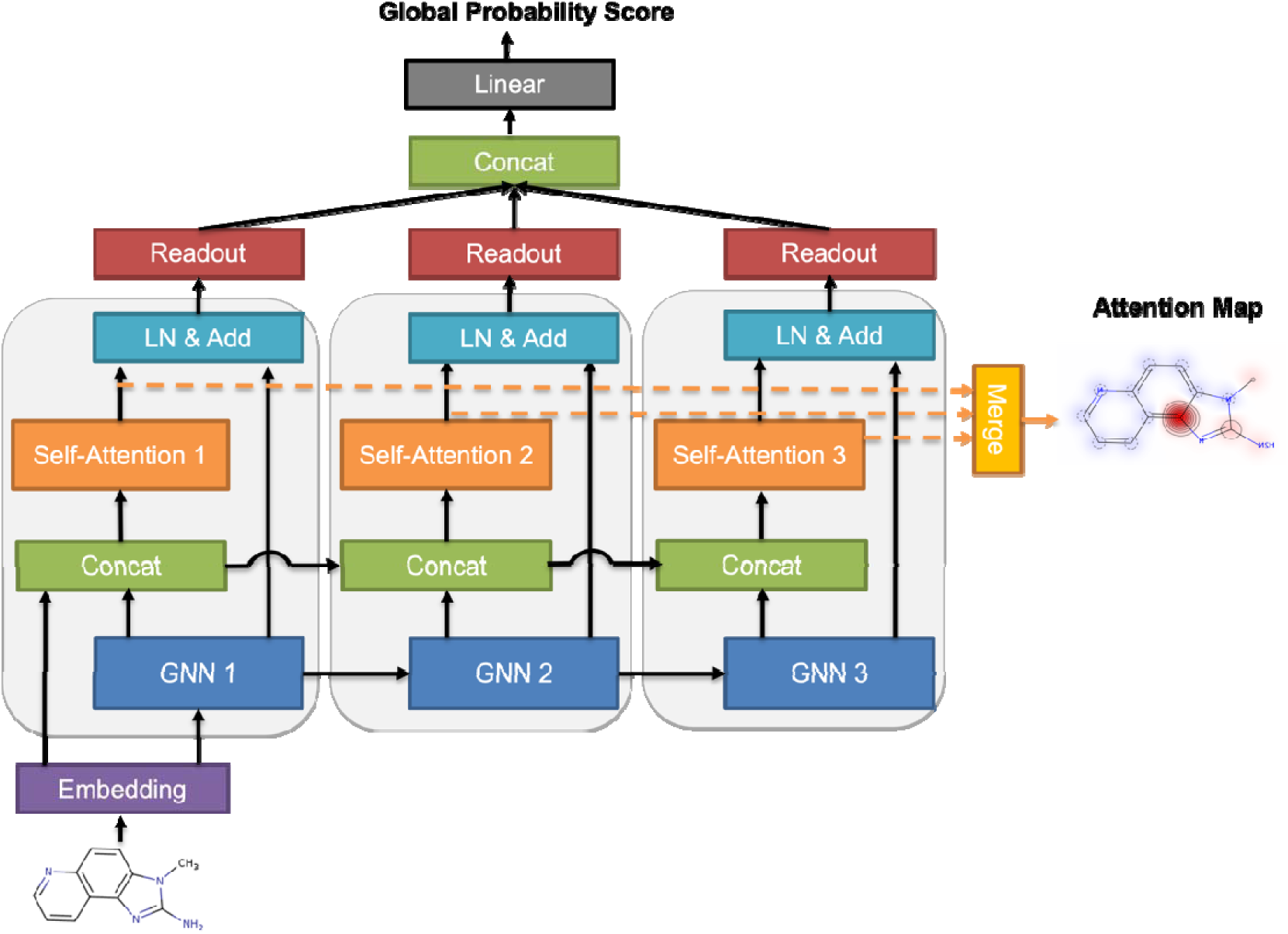
Ligandformer architecture. This is a multi-layer single-head self-attention-based Graph Neural Network framework, for predicting compound chemical property with robust interpretation. Facilitated with visualization technique, the map shows insights on AI model’ rationales on judging which parts of an input molecule impact certain property predictions.

**Fig. S7.**
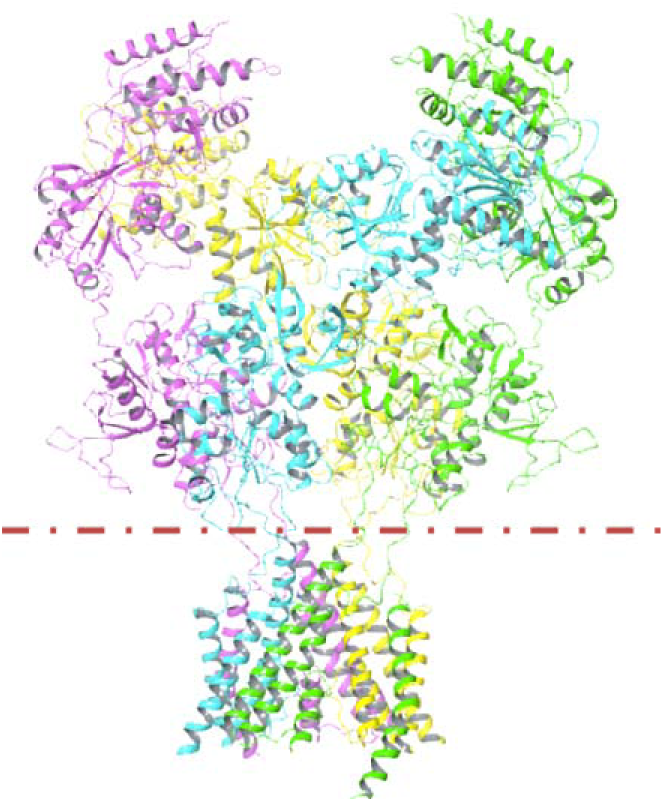
The structure of 7EU7. Only transmembrane region was retained in 100-ns MD simulation (below dotted line.)

**Table S1.** The BBB permeable probability of anti-epileptic drugs in market predicted by Ligandformer model.

| Drug | ID | Probability |
| --- | --- | --- |
| Carbamazepine | DB00564 | 0.9117 |
| Valproic acid | DB00313 | 0.9282 |
| Phenytoin sodium | DB00252 | 0.8325 |
| Oxcarbazepine | DB00776 | 0.94 |
| Diazepam | DB00829 | 0.97 |
| Levetiracetam | DB01202 | 0.95 |
| Fosphenytoin | DB01320 | 0.8445 |
| Flunarizine | DB04841 | 0.9560 |

